# Evaluating passive optical clearing protocols for two-photon deep tissue imaging in adult intact visceral and neuronal organs

**DOI:** 10.1101/018622

**Authors:** R.C. Vlieg, C Gillespie, W.M. Lee

## Abstract

Imaging cellular activities in an entire intact whole organ with light is a grand challenge in optical microscopy. To date, most passive clearing techniques were shown to transform brain, neuronal and embryo tissue into near transparent state for deep tissue imaging. Here, we expand these passive clearing protocols from neuronal tissue (brain and spinal cord) to other visceral organs such as liver and colon and further evaluate their “depth-clearing performance” based on image contrast of endogenous fluorescence structures. We found that SeeDB achieves the highest depth in brain, 3DISCO is adept at clearing liver and spinal cord and ScaleViewA2 in colon. Overall, 3DISCO clears more rapidly than other agents but at a higher cost, while ScaleViewA2 is the most economical however at a slower rate. This study, for the first time, provide a direct evaluation of imaging depth, cost and time amongst passive tissue clearing protocols for different intact organs. In addition, we discuss the possible roles of tissue composition in clearing.

## 1. Introduction

Imaging both the structural and molecular information of intact organs at the cellular level is crucial to the understanding of disease progression and physiology. Modern laser fluorescence microscopy techniques use fluorescent markers that bind to specific cellular proteins so as to track the highlighted cells in intact organs [1]. These laser imaging techniques only operate well in low optical scattering biological samples (cell culture, zebrafish, c. elegans). The key contributing reason behind this is the large optical scattering in bulk tissue that severely limits optical imaging to a shallow depth [2, 3]. Light scattering in tissue can be defined by the transport mean free path (TMFP) [4]. TMFP depends on the scattering coefficient of the tissue that is in turn dependent on wavelength of light and homogeneity of refractive index. On-going efforts to reduce optical scattering (low TMFP) for whole organ imaging includes long wavelength imaging lasers (λ= 1.7 μm), adaptive optics and chemical clearing. Near-infrared pulse lasers emits longer wavelength photons that takes a longer mean free path and hence results in lower scattering coefficients [5]. Adaptive optics (AO) map the scattering coefficient of tissue (transmission matrix) and pre-compensate the incoming laser beam with an optical modulator [6, 7] to directly restore the optical point spread function, thus enabling deep imaging in tissue. Chemical optical clearing methods [8-11] directly replace the interstitial components in tissues with a solution that matches the refractive index of tissue composition (i.e. lipids, n ∼ 1.5), turning intact organs into a state of near transparency.

While the instrumentation-based techniques (imaging laser, adaptive optics) have the advantage of imaging living subjects, they have yet to achieve full imaging depth throughout entire organs [2]. Furthermore, each organ (tissue, colon to brain) are compose differently; connective tissue, interstitial fluids which makes it difficult to compensate without prior knowledge. On the other hand, chemical clearing techniques have achieve imaging depths throughout whole organs (brain, spinal etc) [3] by using agents and equipments that already exist in most biological research facilities. The recent surge in clearing protocols (3DISCO [9], iDISCO [12], SeeDB [8], ScaleView [10], ClearT [13], CUBE[14], PACT [15], Clarity [11] etc) opens up a host of opportunities and increasing popularity amongst researchers for imaging neuronal tissue and embryos [8, 9, 11, 14], but few protocols tackling non-neuronal adult tissue [12, 13, 15]. While new iDISCO [12] and PACT [15] showed significant progress in clearing other organs, their drawbacks are in the non-preservation of endogenous fluorescence. In general, all clearing protocols mainly differ [16] from each other in terms of reagents, cost, clearing time, protocol complexity and tissue optical transparency. Hence, it is unlikely that any single protocol is suitable for all experimental needs. Furthermore, the clearing of some rare diseased samples needs to be done at a calculated risk. To date, there has not been a quantitative study performed to compare the performance, cost and time efficiency in clearing with different protocols [16].

In this paper, we addressed, for the first time, these practical concerns and compared the imaging depth of four optical clearing protocols (glycerol, SeeDB, SCALE, 3DISCO) across four organs (brain, spinal cord, colon and liver) on a high resolution multiphoton microscope. The clearing protocols were chosen because they are passive methods made with commercially available reagents and equipment. We then conducted an assessment of the imaging depth of each optical clearing agent of image contrast using endogenous tissue fluorescence signals and root mean square. The imaging depth in each tissue was compared against the efficiency of each protocol with cost and incubation time. This comparison shall serves as a practical guide to researchers who plan to adopt passive chemical clearing.

## 2. Results

### 2.1. Imaging and contrast

Throughout the imaging sessions, the wavelength of the laser and other imaging parameters (photomultiplier tube: voltage, gain, offset) were kept constant. The excitation wavelength of 900 nm was chosen to excite green fluorescence protein (GFP) excitation and second harmonic generation (SHG) imaging. The imaging channels used to collect the signal include blue (filter BA410-455nm) for SHG from collagen fibers and green (filter BA495-540nm) for endogenous fluorescence and GPF signal. Laser power was gradually increased to compensate for absorption loss along the imaging depth. The step size between each frame in the z-stack is 1 μm. This allowed for relatively fast scanning of the samples, without impairing the resolution of the image greatly. The choice of a 1 μm step size results in a voxel dimension of 1×1×1 μm. In Fig.1, we show, qualitatively, imaging performance using cross sectional views of 3D stacks of each tissue: liver, spinal cord, colon and brain. Control images (PBS solution) are shown in the inset. In non-GFP expressing samples tissue such as liver, auto-fluorescence from sinusoid structures and vasculture are visualized. From Fig.1, it is apparent that the imaging performance have a larger variation for liver and spinal cord, but lower variation for colon and brain. Next we move to quantifying the imaging depth for each tissue sample.

**Figure.1:**
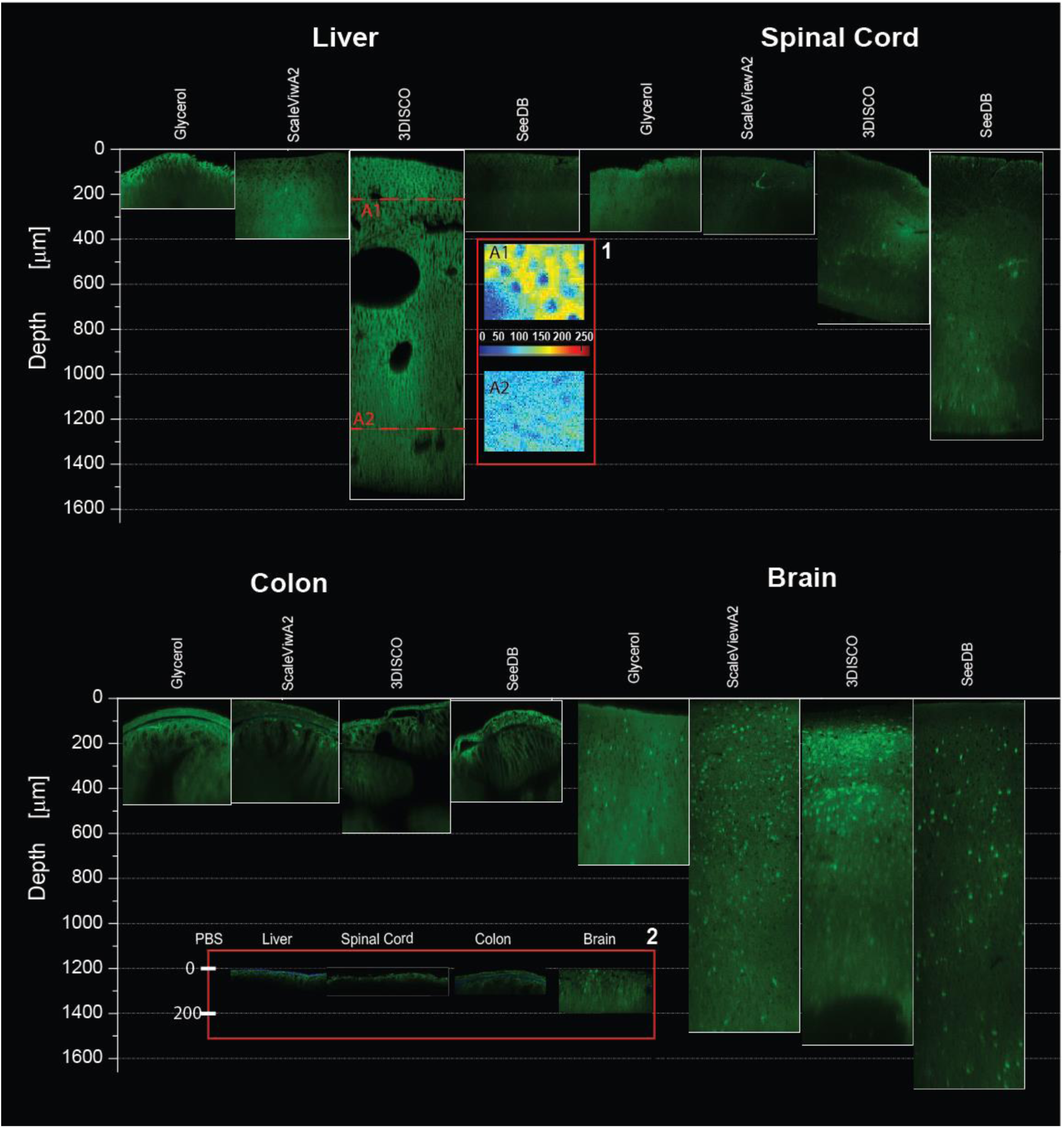
Orthogonal (x-z) slices, taken from multiple z-stacks transverse images, of four tissue types (Liver, Colon, Spinal Cord and Brain) each cleared with four optical clearing techniques (Glycerol, ScaleViewA2, 3DISCO, SeeDB); each z-stack has z interval of 1 μm. Auto-fluorescence image of liver tissue possess characteristic network of vasculatures which serve as negative “contrast” as shown in Inset 1. Inset 2 shows orthogonal slices of control tissue samples immersed in PBS.

### 1.1 Imaging Depth

Previous optical clearing studies use fluorescent particles [8], the optical transmittance[10], endogenous fluorescence[17] and coherence imaging [18] to quantify imaging depth. The full width half intensity maximum (FWHM) of fluorescence cells/particles measures imaging quality at different depths, and endogenous fluorescence (auto-fluorescence) and backscattered light provides imaging of tissues structures (collagen, elastin etc). Auto-fluorescence illuminates distinct structures (vasculature, villi etc) to provide contrast. Table 1 shows the tissue used in this study. In brain tissue, there is an abundant amount of GFP fluorescence cells, but these are sparsely distributed in spinal cord. For other visceral organs where GFP signals are not available, the vasculature in liver and villi distribution in colon serves as form of “negative” contrast. The quantification of contrast here is largely measured by a metric called root mean square (r.m.s) contrast. This is a common approach to measure contrast between objects without relying on spatial frequency content of the image[19] as shown in Eq. 1, where n is the number of pixels and I is the grayscale value.

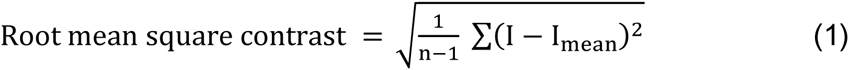

A larger r.m.s contrast also reflects a larger standard deviation of the grayscale values in an image. While this metric is useful for images of no periodicity, the quantification can be sensitive to the variation in mean intensity. This can compromise the reliability of the technique to determine the reliability of contrast in determining depth. We check the reliability of the r.m.s contrast against standard depth imaging with FWHM of fluorescent cells. Brain tissue is used to provide a comparison between the two methods because there are distinct fluorescent cells as well as auto-fluorescence signal. Brain tissue cleared with the four different protocols (Methods). Using 3DISCO protocol, the brain tissue resulted in the transparent amber-glass like appearance similar to the other 3DISCO cleared tissues. SeeDB cleared brain becomes transparent without any colour changes. Tissue volume and optical transparency was increased with ScaleViewA2. On the other hand, brown coloration was observed in the brain tissue with glycerol. The sample images used to quantify imaging depth with the two separate metrics are shown in Fig.2. Fig.2 a i) shows a cropped image of the GFP neurons (false color) that is used to determine contrast (Eq.1) and ii) measured axial full-width half maximum of a GFP fluorescent cell at two depths (160 μm and 1634 um). In Fig.2b, the contrast of each image slice is plotted over the imaging depth and Fig.2c display the full width half maximum (FWHM) of different fluorescence cells is plotted versus the imaging depth. Glycerol and 3DISCO are seen to increase up to 24 ± 14 μm, but SeeDB and ScaleViewA2 showed a more consistent FWHM (16 ± 10 μm). Depth determined with FWHM is given as follows: Glycerol: 934 μm, 3DISCO: 650 μm, SCALE: 1500 μm, SeeDB: 1700 μm) and imaging depth with half of the maximum contrast is given as follows; Glycerol: 346 μm, 3DISCO: 562 μm, SCALE: 691 μm, SeeDB: 934 μm. Imaging depth is decided at the point when contrast value has fallen by half of the maximum contrast (50%) at the tissue surface or that FWHM of fluorescent cells has broaden by more than twice, as shown in Fig. 2a ii). The imaging depths measured using the two metrics are plotted in Fig.2d. The comparison in Fig.2d indicated a greater imaging depth with the FWHM metric than contrast metric (50% contrast and 25% contrast). While the two metric produces similar trend in imaging depth, the absolute value of the imaging depth from contrast varies at different contrast cut off (25% and 50%) from the FWHM measurement. The contrast metric is a suitable metric for comparative measure of imaging depth instead of absolute measure. The contrast metric (50%) is employed to measure the imaging depth in other tissues as shown Fig.3a. 3DISCO is observed to fare much better at clearing liver by over > 1000 μm and moderately in colon by 778 μm and spinal cord with clearing depth of over 700 μm. Clearing liver tissue with glycerol, ScaleViewA2 and SeeDB rendered the appearance of the tissue slightly more white compared to the control sample. 3DISCO hardens liver tissue with an amber coloured glass appearance. None of the protocols were able to fully clear colon tissue over 300 μm which could be explained by tissue composition. Fig.3b shows the 3D reconstruction of tissues with the highest imaging depth in various tissues with brain using SeeDB whereas spinal cord, liver and colon with 3DISCO.

**Figure.2:**
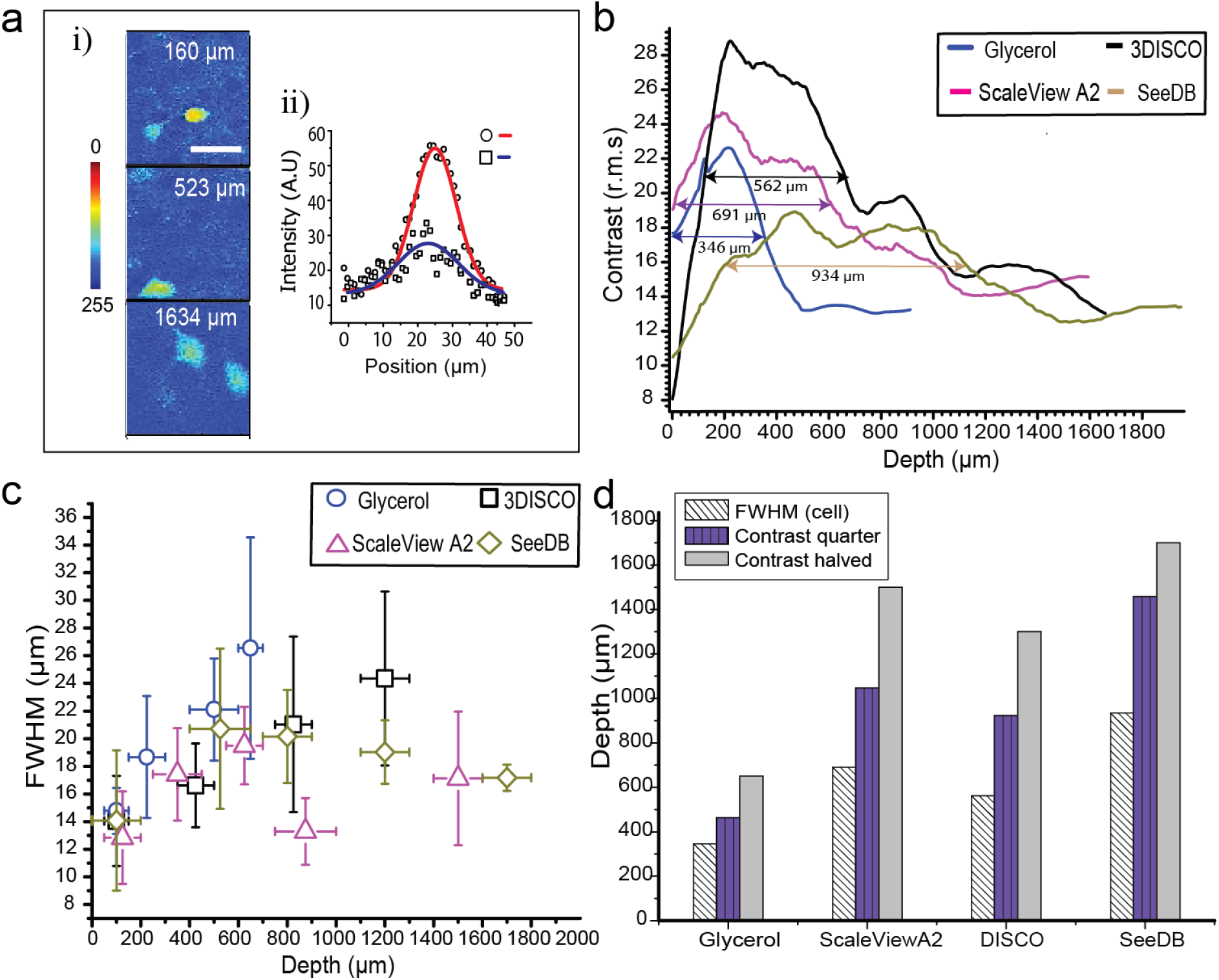
Quantification of imaging depth in cleared brain tissue (SeeDB) with green fluorescence protein positive (GFP+) neurons. (a, i) transverse images at different depths (160 μm, 523 μm and 1634 μm). (a, ii) cross-section intensity plot with Gaussian fits of two GFP+ neurons taken from two different depth (160 μm and 1634 μm). FWHM maximum is derived from the Gaussian fits. (b) Image contrast based on auto-fluorescence signal for brain tissue with different clearing agents. Label for depth determine when contrast drop to 50% from the surface. (c) FWHM of GFP+ cells at different imaging depth with different clearing agents. (d) Comparison of depth determined when the contrast is reduced by 25% (quarter), 50% (half) and FWHM.

**Figure.3:**
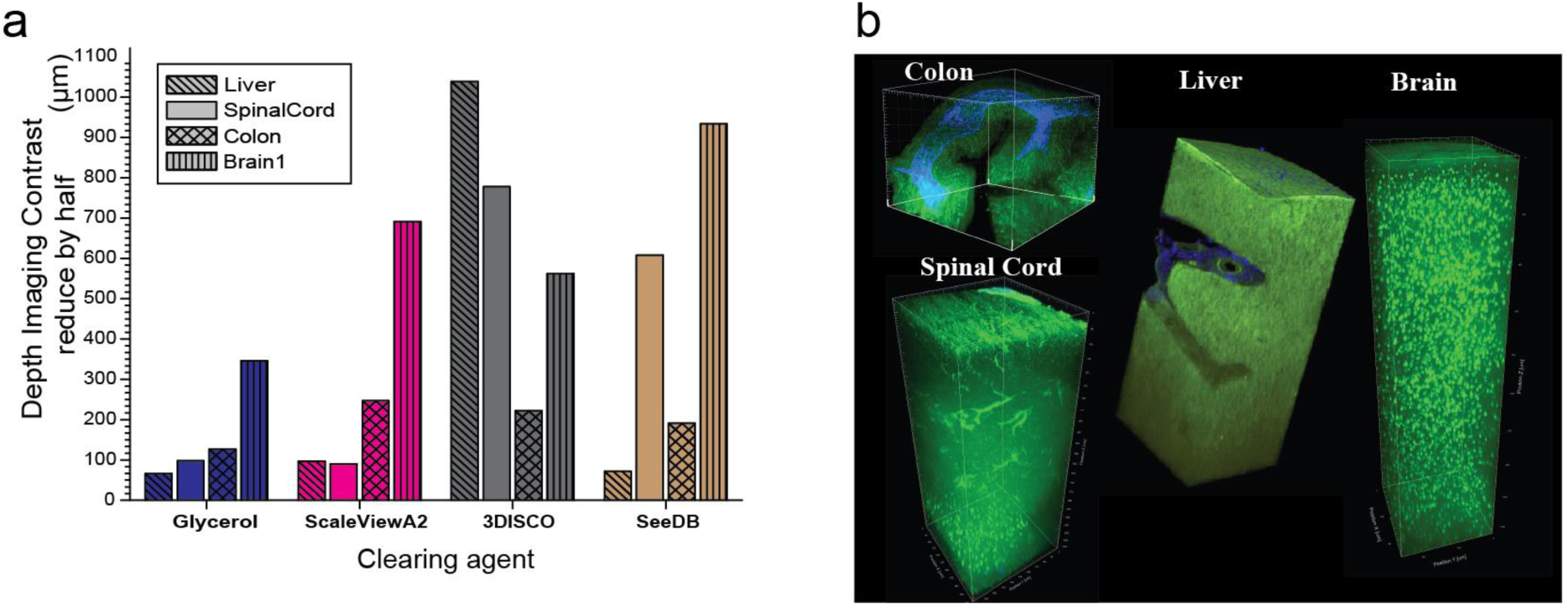
Imaging depth and 3D reconstruction of different cleared tissue defined by contrast (50%). a) plots the imaging depth of each tissue (Liver, spinal cord, colon and brain) for each clearing agent (Glycerol, ScaleviewA2, 3DISCO, SeeDB). b) shows a 3D reconstruction from z-stacks of the tissues with the highest imaging depth

## 3. Discussion

At the heart of optical clearing is the reduction of the scattering coefficient. The clearing process is complex and makes use of mechanisms [3] such as dehydration of tissue, partial replacement of interstitial fluid, molecular structure of clearing agent (hydrophilic/lipophilic) [20] and structural modification of collagen. The efficiency of each clearing process can be further complicated by different organs and tissue types that is difficult to predict experimentally. As the result, the mixture of clearing agents is assessed by their improvement of clearing time, increase optical transparency, compatibility with dyes and cell markers and also its viability to clear different tissue types. Hirshburg *et al* [20] showed that the optical clearing agent with high affinity to form hydrogen bridges to homogenized tissue refractive index correlate to high efficiency in optical clearing than index matching. More recent clearing compounds have targeted lipids to enhance clearing. For example, urea (ScaleView) removes lipids and maintains fluorescence proteins, however compromises the structural integrity of tissue. Hydrogel-based clearing (Clarity, PACT) replaces lipid with monomers that stabilized the tissue structure over a long time, but facilities the addition of proteins through monomers. Every iteration protocol tries to address shortcomings in the previously developed protocols, however, accompanied with drawbacks of its own. Based on the current trend, it is more likely to pick suitable protocols specific to organs instead of a multipurpose, low cost and high efficient clearing agent. In the future, it will be useful to combine live cellular level imaging techniques to study the uptake of new clearing agents at single cell level so as to determine the local effects of clearing agents.

### 1.2. Clearing agents – variation in organs

Out of four passive clearing protocol, we determined the optimal protocol for the adult visceral and neuronal organs. Our imaging results clearly demonstrate a significant variation between different organs. The effect may be attributed to the different tissue layers in each organs. For example, brain is mainly comprise of neuronal networks and small vasculatures that has not collagen. However, in spinal cord, there is a network of connective tissues (collagen) with nerve bundles (myelinated) that increases scattering. In lower colon, the composition is much more complex where there are multiple separate layers of columnar epithelial layers (villi), muscle cells and connective tissues that increases optical scattering. Liver, on the other hand, is highly vascularized (veins, ducts) with densely distributed hepatic cells. The tissue structure alone could influence the efficiency of passive diffusion into interstitial compartments. In fact, our results do demonstrate, to some degree, the influence of tissue structure to clearing through different organs. Brain and colon display a sharp contrast in optical clearing with all four clearing protocol. This could be explained by the difference in tissue composition where brain possess a high level of homogeneity than colon. On the other hand, liver and brain have similar tissue composition; cells (hepatic and neuronal cells) and high density of vasculatures, aside from sparsely distribution collagen fibers in liver. However, liver possess larger diameter vascular networks than brain and other tissue. Our repeatable observation that 3DISCO showed excellent clearing performance above other clearing agents. This suggests that larger vascular tissue requires rapid dehydration and replacement of index matching fluid. Next, myelinated spinal tissue (high lipid content) increases optical scattering. Our results showed that both dehydration (3DISCO) and high osomatic pressure (SeeDB) promotes improved clearing in myelinated tissue as oppose to lipid disruption by urea in ScaleView. This is consistent with previous clearing experiments [14]. In the future, more efforts will be needed to look into the roles of diffusion/uptake rate of clearing agent with varying viscosity and molecular weight in the complex layers of the lower colon tissue.

### 1.3. Time and cost

The primary objective of this work was to compare the performance (depth, cost and time) of the four optical clearing agents on a standard multiphoton microscope platform and also assess imaging depth for different tissues with endogenous fluorescence signal. A drawback with relying on the intrinsic fluorescent structure of the tissue is the influence of background signal. For more homogenous sample, such as brain, the cells or vasculature needs to be highlighted with fluorescent markers and dyes. The advantage of auto-fluorescence technique is in the flexibility of quantifying non-labeled sample and does not to rely on distribution of cells and undesirable photobleaching effects. Apart from the depth quantification, clear tissue still poses significant challenge to achieve diffraction limited imaging at deep depth. The reasons include limited working distance of existing microscope objective lenses and increasing spherical aberrations. There are ways to overcome this limitation, such as manipulating the refractive index of the objective immersion medium or by using special long-working distance microscope objectives [21]. These optical approaches will become more prevalent as more researchers adopt clearing techniques.

To conclude, we provide the analysis of both cost and time for different clearing agent and different tissue. In Fig.4, we plotted the depth achieved with respect to cost and incubation time. Fig.4a shows that ScaleViewA2 is the most economical but at the much slower clearing rate. On average, 3DISCO cleared all the organs more rapidly but it was more costly. For brain tissue, glycerol achieved a relatively fast clearing time while imaging depth stayed restricted. Glycerol was found to work nicely at low cost and reasonably fast in clearing depth in brain. Our result (unpublished) also verifies that glycerol is aggressive (dissolves collagen) and lowers GFP fluorescence due to quenching. None of the clearing techniques attempted here can clear entire colon tissue for high resolution imaging. 3DISCO achieved the deepest imaging depth, which makes it the best option for deep-tissue imaging of colon tissue. It is also good to note that clearing colon tissue with glycerol only took 20 minutes (∼254 μm/h optical clearing speed). In summary, our results indicated that 3DISCO is suitable for time-constrained users, and SeeDB is suitable for users requiring an increased imaging depth at a greater cost. Both ScaleView and Glycerol are cost efficient with Scaleview achieving a deeper imaging depth in brain. This study provides sufficient information for any user to embark on the appropriate clearing protocol for their needs.

**Figure.4:**
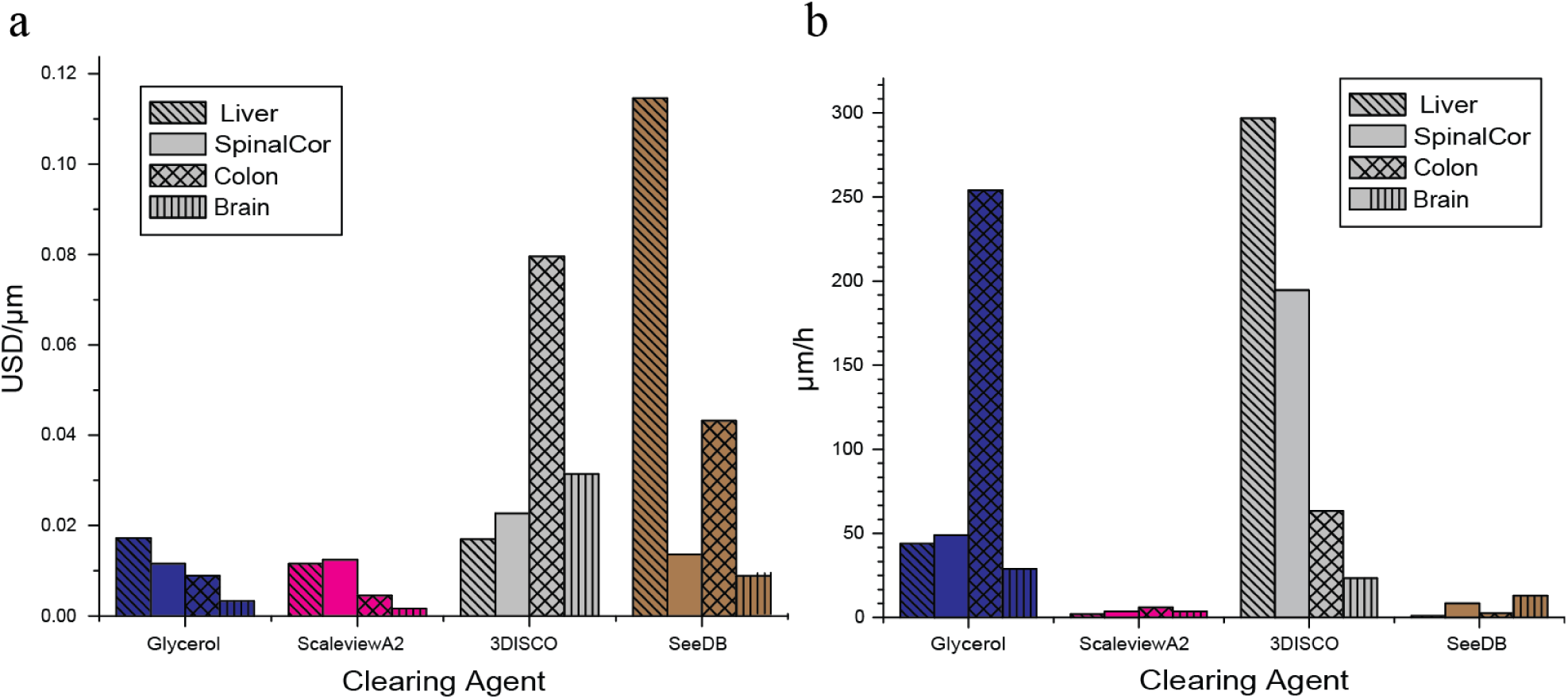
Column plots of two parameters (cost -USD and time – hours) relative to the achieved imaging depth achieved in Fig.3c. a) Cost is calculated by adding all the cost for each ingredient used in a particular clearing agent for a single tissue sample. The total cost is then divided by the imaging depth. b) show the rate of clearing for different clearing agent based on imaging depth per incubation time.

## 2. Methods

### 2.1. Multiphoton Microscope and sample chamber

A standard multiphoton (Olympus© FVMPE-RS) microscope, with 2 line lasers (a tunable 680-1300 nm line and a fixed 1040 nm line - Spectra Physics DeepSee InSight laser), was used to image the samples. While endogenous fluorescence signal (auto-fluorescence) shows structural details across the whole organ after clearing, there is still considerable loss of fluorescence signal due to limited working distance of the objective lens, optical absorption, and spherical aberrations. A standard working distance objective; Olympus© water immersion XLPlan N objective (25x magnification, numerical aperture: 1.05, working distance: 2.0 mm) is used in imaging, as illustrated in Fig.5a. With longer working distance objectives, it is possible to improve the imaging quality over the imaging depth [10]. While absorption can be compensated with increasing laser power, the amount of spherical aberration can be minimized by matching refractive index between the sample and objective lenses. A custom-made chamber is used to immense organ in clearing solution for minimal refractive index variation between coverslip and organ. The sample chamber is made out of (polydimethylsiloxane, PDMS) and molded from 3D printed mold Fig.5b. Silicone grease is applied onto a glass cover slide as a temporary seal. Spherical aberrations between the objective lens and the coverslip are corrected by adjusting the collection collar. Collar correction is used to correct for additional first order spherical aberrations and defocusing [22]. The collar correction was corrected for the surface of the tissue for all imaging to ensure clear comparison of the imaging performance. A mold of the chamber was first designed in SolidWorks© and 3D printed (UpMINI 3D) as shown in Fig.5b. The negative mold is filled with a silicone (polymethylsiloxane, PDMS) solution and once the silicone solution has become firm, it is removed from the mold. Fig.5b shows the mold and the silicone chamber. For samples of varying sizes, different sized imaging chambers are used, ranging from 3 mm in height for spinal cord tissue up to 10 mm for half mouse brain samples. For larger tissue (e.g. liver and brain), the bottom of the chamber is coated with dental cement (Paladur, Pala) to seal the chamber. To make the dental cement, the two components of Paladur, powder and liquid, are mixed together in a 10:6 (5:3) ratio, cause a polymerization reaction, which create a viscous fluid. This fluid hardens in an approximately 3 minute time window, giving the user time to mold the cement in the desired shape. Fig.5 c and d shows the 3D reconstruction and cross-sectional slice of whole brain tissue (GAD67-GFP knock-in mouse) cleared with SeeDB respectively.

**Figure.5.**
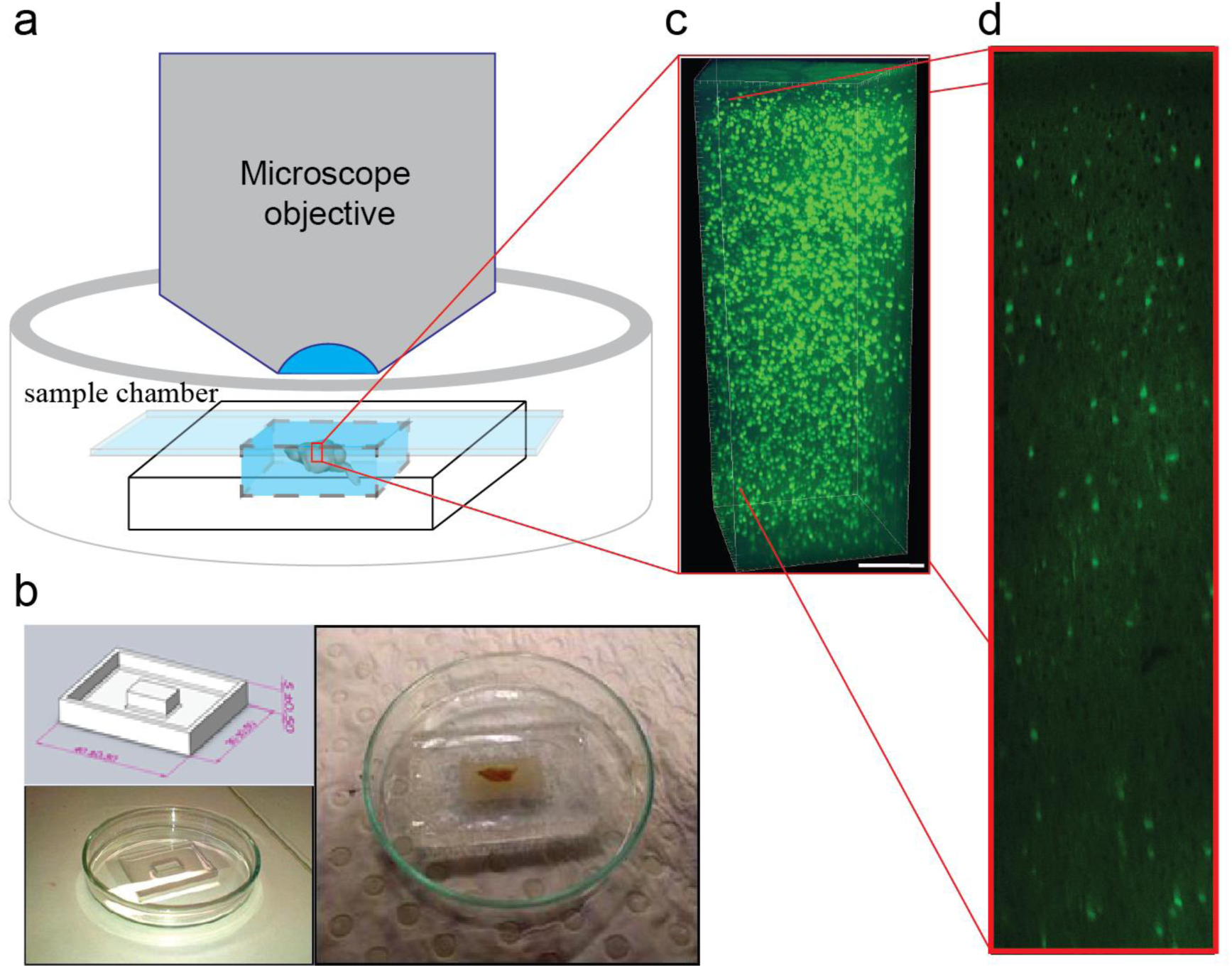
– Multiphoton imaging of cleared organs in customized sample chamber. (a) a typical imaging session where the organ is immersed in the clearing agent within an enclosed silicone chamber sealed with vacuum grease and a glass coverslip. A drop of water is placed on top of the coverslip to match the immersion medium of the objective. (b) (Left-top) shows a 3D printed ABS negative mold of the sample chamber. (Left - bottom) Silicone is made by curing polydimethylsiloxane (PDMS) under 70°C for 20 minutes. (Right) shows a piece of a half mouse brain placed within the chamber before imaging. (c) 3D reconstructions partial brain organ after clearing with SeeDB. (d) shows an axial slice of the tissue showing individually GFP+ neuronal for over 1.8 mm.

### 2.2. Clearing agent

In Table 1, we list the protocols used here, namely, glycerol, SCALE, SeeDB, 3DISCO. Glycerol is chosen as our reference clearing agent because it is considered as one of the most basic clearing procedures available.

**Table.1.** Clearing Agent and sample origin

| Clearing Agent | Organ | Mice |
| --- | --- | --- |
| Glycerol,<br>3DISCO<br>ScaleViewA2,<br>SeeDB | Liver | C57Bl/6 ubiquitin-GFP mouse<br>with adoptively transferred OT-1 GFP cells |
|  | Brain, gut<br>and spinal<br>cord | GAD67-GFP knock-in mouse |

### 2.3. Sample preparation

Samples for the experiments were:

In order to reduce variability in our samples, the organs are prepared in the following manner;

1. Mice were anesthetized using chloral hydrate (400mg/kg) or ketamine (10mg/kg)/xylazine (10mg/kg).
2. They were perfused with 4% saline for 5-10 min followed by 5 min of 4% (wt/vol) paraformaldehyde (PFA) in 0.1 M PB, using a syringe operated manually.
3. Organs of interest were dissected and post-fixed by immersion using 4% (wt/vol) PFA overnight.
4. Organ was placed in a Petri dish with PBS and extra tissue was removed and/or sliced into required sample size.
5. Samples was immersed in PBS and stored at 4 °C in the fridge.

### 2.4. Clearing protocols

Each protocol consists of basically three main steps: clearing agent preparation, sample preparation and incubation of the sample in the clearing agent; and preparation of the cleared sample for imaging. The preparation steps together with the incubation period will take up mostly of the time The protocols used in this study are based on the published protocols for SeeDB, 3DISCO and Scale [8,9,10]. As the equipment and resources are slightly different than that described in these papers, the protocols used here are altered according to the available laboratory equipment.

#### 2.4.1. Glycerol

1. Prepare 80% glycerol/20% PBS solution.

##### 2.4.1.1. Tissue clearing

1. Incubate sample in 20 ml glass vial for ∼1 hours in 80% glycerol solution
2. Put sample in fresh 80% glycerol solution in the imaging chamber.
3. Put imaging chamber under the microscope and acquire images.

#### 2.4.2. 3DISCO

1. Prepare 50%, 70%, 80%, 100% tetrahydroforan (THF) working solutions from the stock solution in 20 ml glass vials, dilute with distilled water. Mix by shaking for few minutes.
2. Solutions should be stored in the dark, tightly sealed, at room temperature. No more than two weeks
3. Prepare dibenzyl ether (DBE) solution, 100 ml into small glass bottle, no dilution needed (99-100%).
4. DBE should be stored in the dark, tightly sealed at room temperature. Storage for months to years can lead to peroxides in the solution which are explosive.
5. Prepare DCM solution, 100 ml into small glass bottle, no dilution needed (99-100%).
6. Store in the dark, tightly sealed at room temperature. For best results, do not use for longer than two weeks.

##### 2.4.2.1. Tissue clearing

1. Transfer the sample directly into the 20 ml vial containing the clearing solution in accordance with Table 2.
2. Add clearing solutions according to clearing steps described in Table 2 using a Pasteur pipette.
3. Close the lid of the glass vial and place on the turning wheel and cover the sample with aluminium foil.
4. Rotate the sample at constant speed (20-40 r.p.m.)
5. When appropriate time has passed, place the sample in the next clearing solution.
6. Repeat step 5-8 until last clearing solution is reached (DBE). The incubation time for DBE is important: 15-20 min for spinal cord tissue; larger organs from 6 hours to overnight for adult mouse brain.
7. Proceed to image as soon as the tissue is cleared. DBE degrades the fluorescence signal; waiting one day already decreases fluorescence.
8. Once the tissue is cleared, imaging should commence soon after. Leaving the sample overnight in the DBE solution reduces fluorescence signal and tissue is less clear as well. Once the tissue is imaged, it can be discarded.

**Table 2.** 3DISCO immersion protocol

| Reagent | Spinal cord, liver, colon | Brain |
| --- | --- | --- |
| 50% (vol/vol) THF | 30 min | 1 h |
| 70% (vol/vol) THF | 30 min | 1 h |
| 80% (vol/vol) THF | 30 min | 1 h |
| 100% (vol/vol) THF | 3 x 30 min | 1h, overnight, 1h |
| DCM | 20 min | - |
| DBE | $\geq 15$ min | $\geq 3$ h |

#### 2.4.3. ScaleViewA2 [10] (available from Olympus)

1. Dissolve 240.24 g of urea crystals
2. Add 10 ml of 10% (wt/vol) Triton X-100 solution.
3. Add 100 g of glycerol
4. Mix well by stirring.
5. Add Milli-Q water to make 1,000 ml and stir until well mixed.
6. Store at room temperature.

##### 2.4.3.1. Tissue clearing

1. Transfer the sample into ScaleViewA2 solution (20 ml/0.5 g tissue) in 20 ml glass vial.
  - Salt from the PBS fixation solution may cause white precipitates. When white precipitates are observed, incubation solution should be refreshed with new ScaleViewA2 solution.
2. Incubate the sample for 2-14 days at 4 °C or room temperature with gentle shaking. Exchange ScaleViewA2 if necessary (see bulletpoint at Step 1). Observe the sample regularly, to assess the need for fresh ScaleViewA2 solution or the stadium of optical clearing the sample is in.
  - Older mice (3 > weeks old) may introduce some issue during clearing with ScaleViewA2 solution. Spitting brain in half or introducing a few slits may prove beneficial for clearing of the brain.
3. After observing satisfactory clearing of the sample, place the sample in the imaging chamber immersed in fresh ScaleViewA2 solution.
4. Image the sample under the multi-photon microscope.
5. Sample can be stored at 4 °C, or room temperature, in fresh ScaleViewA2 solution. Fluorescence is not lost over time.

#### 2.4.4. SeeDB

1. Prepare the fructose 20% (w/v), 40% (w/v), 60% (w/v), 80% (w/v), 100% (w/v), SeeDB, by adding distilled water to the fructose, see Table 3. α-thioglycerol (THF) is added just before tissue incubation. When solution is heated to 50 °C, it should be cooled down to room temperature when adding α-thioglycerol. To dissolve the fructose solution in the 100% (w/v) and the SeeDB solution, place the solution at 50 °C using the oven in 50 ml conical centrifuge tubes. Shake the solutions for couple of minutes to completely dissolve the fructose. SeeDB is prepared on weight/weight ratio; 80.2% (wt/wt). The other fructose solutions are prepared on weight/volume ratio. See Table 3 for the quantities in which the different chemicals should be mixed to obtain the needed clearing agents.
2. Solution should be stored for no longer than 7 days; all solutions should be freshly prepared before clearing of tissue. Store at room temperature.

**Table 3:**
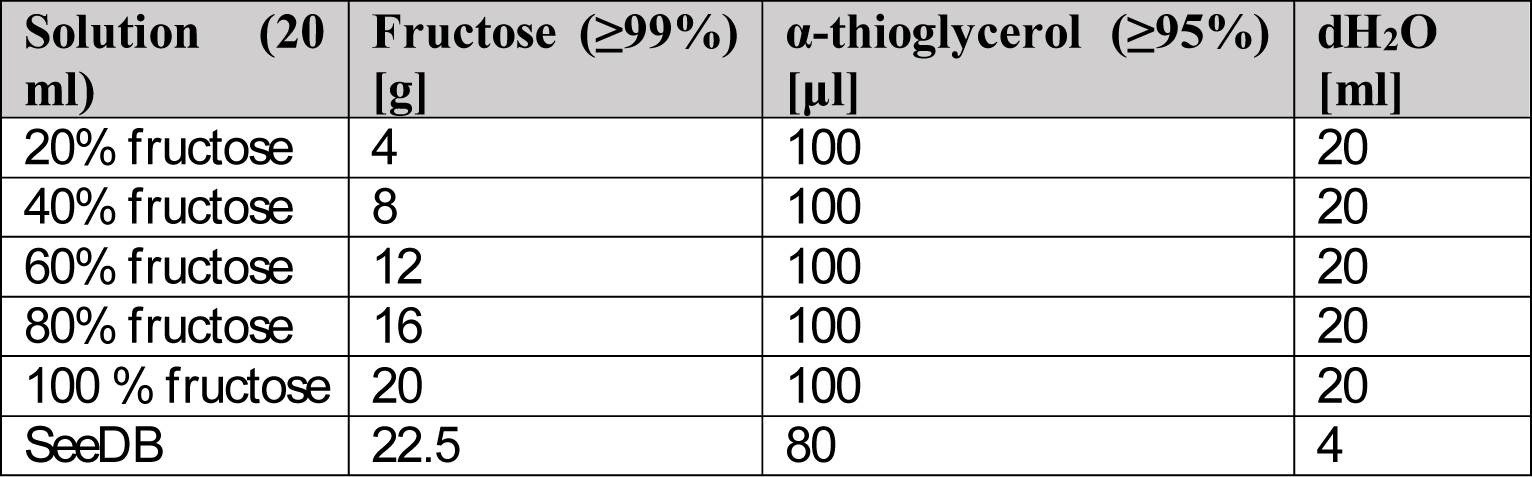
Solutions required for the SeeDB protocol and their composition

| **Solution (20 ml)** | **Fructose ($\geq 99\%$) [g]** | **α-thioglycerol ($\geq 95\%$) [μl]** | **dH<sub>2</sub>O [ml]** |
| --- | --- | --- | --- |
| 20% fructose | 4 | 100 | 20 |
| 40% fructose | 8 | 100 | 20 |
| 60% fructose | 12 | 100 | 20 |
| 80% fructose | 16 | 100 | 20 |
| 100 % fructose | 20 | 100 | 20 |
| SeeDB | 22.5 | 80 | 4 |

##### 2.4.4.1. Tissue clearing

1. Transfer the sample to a 50 ml conical tube containing 20 ml of 20% (w/v) fructose solution.
2. Place the conical tube on a tube rotator for 4 to 8 hours (depending on the sample) at constant speed of 4 r.p.m. at room temperature.
3. Place the sample in a new 50 ml conical tube containing 20 ml of the next fructose solution. Place tube in the tube rotator and incubate the sample for the appropriate time, see Table 4.
4. Repeat Step 4 until the sample has been incubated in the last immersion step for the appropriate amount of time.
5. Put the cleared sample in the imaging chamber together with SeeDB solution.
6. Using the SeeDB solution in the chamber will maintain the same refractive index.
7. Place the sample under the microscope.

**Table 4:** SeeDB immersion protocol

| **Immersion solution** | **20% fructose** | **40 % fructose** | **60% fructose** | **100% fructose** | **SeeD B** |
| --- | --- | --- | --- | --- | --- |
| **Incubation time** | 8 h | 8 h | 8 h | 12 h | 24 h |

## 3. Contributions

R.C.V and W.M.L worked together on the planning and execution of the experiment. R.C.V conducted the analysis and quantification of the data with input from W.M.L. C.G. provided technical support in clearing and imaging. All authors discussed the results. R.C.V and W.M.L wrote the manuscript.

## 4. Acknowledgements

We thank Prof. John Bekkers, Dr Norimitsu Suzuki and Assoc Prof Ian Cockburn for providing the samples for this study. Mr Lyle Halliday for 3D reconstruction of liver in Fig.3b. Dr Clemens Alt for providing critical comments on the manuscript.

